# The McGill Face Database: validation and insights into the recognition of facial expressions of complex mental states

**DOI:** 10.1101/586453

**Authors:** Gunnar Schmidtmann, Ben J. Jennings, Dasha A. Sandra, Jordan Pollock, Ian Gold

## Abstract

Current databases of facial expressions of mental states typically represent only a small subset of expressions, usually covering the basic emotions (fear, disgust, surprise, happiness, sadness, and anger). To overcome these limitations, we introduce a new database of pictures of facial expressions reflecting the richness of mental states. 93 expressions of mental states were interpreted by two professional actors and high-quality pictures were taken under controlled conditions in front and side view. The database was validated with two different experiments (N=65). First, a four-alternative forced choice paradigm was employed to test the ability of participants to correctly select a term associated with each expression. In a second experiment, we employed a paradigm that did not rely on any semantic information. The task was to locate each face within a two-dimensional space of valence and arousal (mental state – space) employing a “point-and-click” paradigm. Results from both experiments demonstrate that subjects can reliably recognize a great diversity of emotional states from facial expressions. Interestingly, while subjects’ performance was better for front view images, the advantage over the side view was not dramatic. To our knowledge, this is the first demonstration of the high degree of accuracy human viewers exhibit when identifying complex mental states from only partially visible facial features. The McGill Face Database provides a wide range of facial expressions that can be linked to mental state terms and can be accurately characterized in terms of arousal and valence.

## Introduction

Faces represent a special, very complex class of visual stimuli and have been extensively studied in a wide range of research areas. In particular, facial expressions are among the most important sources of information about the mental states of others. The capacity to make mental state inferences, whether from faces or other sources, is known as Theory of Mind (ToM), and it is widely agreed that this capacity is essential to human social behavior. There is also substantial evidence that a ToM deficit may be associated with a variety of clinical conditions, notably autism (Baron-Cohen et al. (1997, 2001)) and schizophrenia (Bora et al. (2009), Brüne (2005), Harrington et al. (2005), Sprong et al. (2007)). Hence, the assessment of ToM is important for the exploration of social cognition in healthy individuals as well as in some patients. It may also be useful to measure a change in the social capacities of patients in psychotherapy. The “Reading the Mind in the Eyes” Test (Baron-Cohen et al. (1997, 2001)) is a common ToM test in which participants have to choose a mental state term that best characterizes the expression in a picture of someone’s eyes. However, only a small proportion of possible mental states are tested, and the stimuli themselves are of inconsistent quality with respect to image resolution, luminance and perspective. Most other comparable databases of facial expressions of mental states typically only include a small subset of expressions, typically the basic emotions proposed by Paul Ekman (e.g. Ekman, 1992): fear, disgust, surprise, happiness, sadness, and anger) – the emotional expressions that are considered universal. However, multiple secondary emotions where two or more primary emotions are mixed (e.g. hatred being a mix of anger and disgust, are highly under-represented in the databases available. One exception is the “Mind Reading” database (DVD, Baron-Cohen et al., 2004 that contains a much wider range of mental states. The Mind Reading DVD is computer-based platform developed to help individuals diagnosed along the autism spectrum to recognize facial expressions. It contains 412 mental state concepts, each assigned to one of 24 mental state classes. However, it is designed for commercial and clinical use and specifically targets patients with autism spectrum disorder and Asperger syndrome. A list of popular face stimuli databases is shown in Table 1. Most databases only represent a very small subset of emotions encountered in daily life and often in exaggerated form. To overcome these limitations, we have developed and validated a large new database of pictures of facial expressions – the McGill Face Database – that reflects some of the richness of human mental states. The database contains high-resolution pictures of 93 expressions of mental states that were interpreted by two professional actors (one male and one female) in front and side view – 372 images in total. In this paper, we present two different experiments to investigate subjects’ ability to recognize the facial expressions in the Database. In experiment 1, we employ a four-alternative forced choice paradigm, based on previous studies (Baron-Cohen et al. (1997, 2001)). The task for the observer in this experiment was to choose, out of four terms, the one that best identifies the mental state expressed. Given that a particular “correct” term is only a representation of the actors’ interpretations of the mental state, a second validation experiment (experiment 2) was carried out, which did not rely on the semantics of the mental state terms. Instead, the observers located each face within a two-dimensional space of valence and arousal (mental state – space) employing a “point-and-click” paradigm (Jennings et al. (2017)).

**Table 1.**
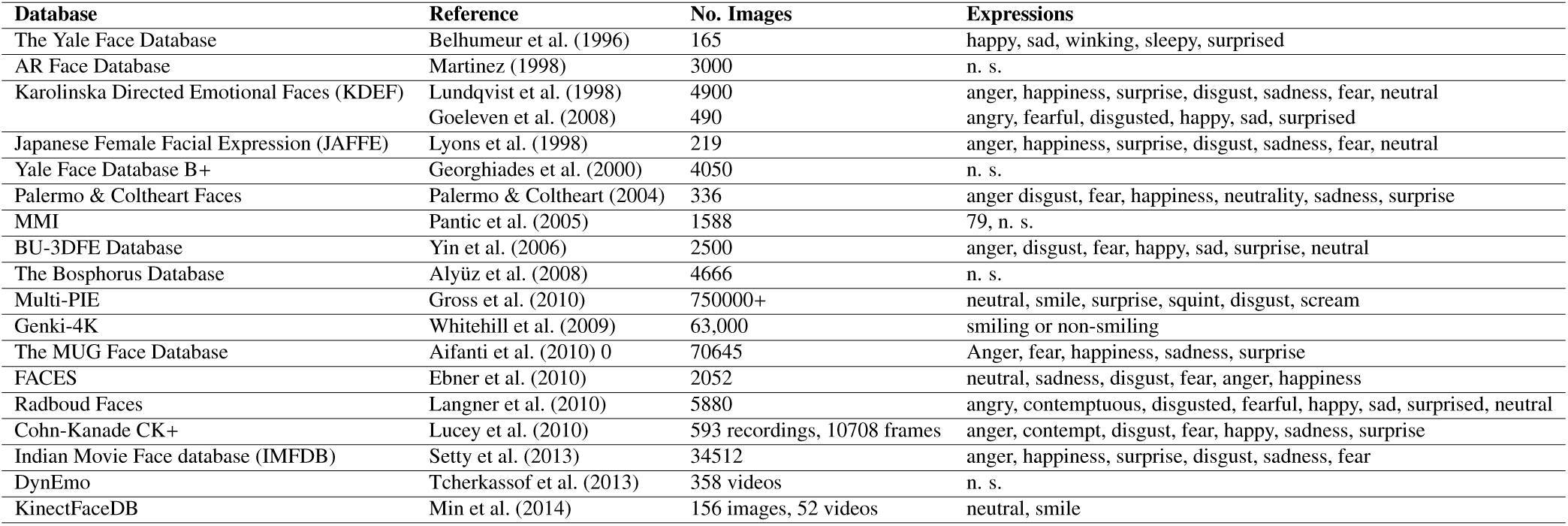
Summary of face databases (n.s.: not specified)

## Database

### Actor Recruitment

Five male and five female professional native English-speaking actors were invited to take part in an audition. The actors’ performance was judged by a panel of two of the authors and a theater-experienced Professor of Drama and Theatre in the McGill Department of English. During the audition, one male and one female actor engaged in various improvisation exercises. The “best actors” were those who exhibited the most precise, nuanced, and yet readable range of emotional expression in their faces, i.e. that clarity of emotional expression – as captured by the camera - was paramount. Some actors were better able to convey different emotions through subtle recalibration of facial expression while others either got “stuck in look” or fell into exaggerated or melodramatic countenances. The two best-performing actors (male, age 29, female, age 23) were chosen to take part in a photo shoot based on a majority vote. The actors gave informed consent and signed an agreement allowing for the pictures to be used for research and other non-commercial purposes. The actors were compensated for their work.

### Images

#### Equipment

The pictures were taken by a professional photographer with a Canon 70D digital camera mounted on a tripod at a distance of 1.5 m from the actor. The optic was a Canon 85 mm, f1.8 with a shutter speed of 1/60th and an aperture of f5.6 and a sensitivity of ISO 100. Two separate flashes—a Canon 580 EX and a Canon 430 EXII (both set with exposure compensation at +1) were placed at the appropriate distance. One of the flashes had a reflector umbrella.

#### Image Acquisition

The pictures were taken in two separate sessions at a studio specifically prepared for that purpose. During the sessions, the actor was positioned in front of a white screen. The instructor provided the mental state term and read the corresponding short explication provided in the Glossary in Appendix B of Baron-Cohen et al. (2001). The actor was given as much time as needed to prepare the interpretation for the relevant expression. When the actor gave a hand signal to the photographer, a single picture was taken in front view. Importantly, in order to guarantee a natural interpretation of a given expression, we did not restrict the head tilt. The actor then immediately turned to face a mark 30° from the camera, and a second picture was taken. This procedure was repeated three to four times for each of 93 mental state terms used in the Reading the Mind in the Eyes Test (Baron-Cohen et al. (2001)) (Table 2 in the Appendix).

**Table 2.** Summary of terms in the McGill Face Database

|  | English |  | English |
| --- | --- | --- | --- |
| 1 | Accusing | 48 | Grateful |
| 2 | Affectionate | 49 | Guilty |
| 3 | Aghast | 50 | Hateful |
| 4 | Alarmed | 51 | Hopeful |
| 5 | Amused | 52 | Horrified |
| 6 | Annoyed | 53 | Hostile |
| 7 | Anticipating | 54 | Impatient |
| 8 | Anxious | 55 | Imploring |
| 9 | Apologetic | 56 | Incredulous |
| 10 | Arrogant | 57 | Indecisive |
| 11 | Ashamed | 58 | Indifferent |
| 12 | Assertive | 59 | Insisting |
| 13 | Baffled | 60 | Insulting |
| 14 | Bewildered | 61 | Interested |
| 15 | Cautious | 62 | Intrigued |
| 16 | Comforting | 63 | Irritated |
| 17 | Concerned | 64 | Jealous |
| 18 | Confident | 65 | Joking |
| 19 | Confused | 66 | Nervous |
| 20 | Contemplative | 67 | Offended |
| 21 | Contented | 68 | Panicked |
| 22 | Convinced | 69 | Pensive |
| 23 | Curious | 70 | Perplexed |
| 24 | Deciding | 71 | Playful |
| 25 | Decisive | 72 | Preoccupied |
| 26 | Defiant | 73 | Puzzled |
| 27 | Depressed | 74 | Reassuring |
| 28 | Desire | 75 | Reflective |
| 29 | Despondent | 76 | Regretful |
| 30 | Disappointed | 77 | Relaxed |
| 31 | Dispirited | 78 | Relieved |
| 32 | Distrustful | 79 | Resentful |
| 33 | Dominant | 80 | Sarcastic |
| 34 | Doubtful | 81 | Satisfied |
| 35 | Dubious | 82 | Serious |
| 36 | Eager | 83 | Skeptical |
| 37 | Earnest | 84 | Stern |
| 38 | Embarrassed | 85 | Suspicious |
| 39 | Encouraging | 86 | Sympathetic |
| 40 | Entertained | 87 | Tentative |
| 41 | Enthused | 88 | Terrified |
| 42 | Fantasizing | 89 | Thoughtful |
| 43 | Fascinated | 90 | Threatening |
| 44 | Fearful | 91 | Uneasy |
| 45 | Flirtatious | 92 | Upset |
| 46 | Flustered | 93 | Worried |
| 47 | Friendly |  |  |

#### Image Selection

A focus group, consisting of six referees (four females and two males) were presented with the different images for a given expression and asked to compare their quality and expressivity of mental state. Four out of six referees had to agree on a picture for it to be selected for inclusion in the database. The full database can be downloaded at: McGill Face Database.

#### Image Specificities

The database contains 372 jpegimage files with a resolution of 5472 × 3648 pixel (colour space profile: sRGB IEC61966-2.1). The size of each image is 7.3 MB. The image files have not been post-processed. Raw image files are available upon request from the first author.

## Experiment 1

### Methods

#### Subjects

All participants were recruited via the McGill Psychology Human Participant Pool or via public advertisements. 33 individuals (7 males, 26 females, mean age 21 years, ± 2.96 SD) participated in Experiment 1. All subjects were native English speakers and were naïve as to the purpose of the study. Subjects had normal or corrected-to-normal visual acuity. Informed consent was obtained from each observer. All experiments were approved by the McGill University Ethics committee and were conducted in accordance with the original Declaration of Helsinki.

#### Apparatus

The face stimuli were presented using MATLAB (MATLAB R 2016b, MathWorks) on either a CRT monitor running with a resolution of 1600 × 1200 pixel and a frame rate of 60 Hz (mean luminance 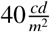) under the control of an PC (3.2GHz) or on a MacBook Pro (2015, 3.1 GHz) with a monitor resolution of 2560 × 1600 pixel. The viewing distance was adjusted to guarantee an equal image size of 20.91° × 13.95° on both systems. Experiments were performed in a dimly illuminated room. Routines from the Psychtoolbox-3 were employed to present the stimuli (Brainard & Vision (1997)).

#### Procedure

A four-alternative forced choice paradigm was employed to test the ability of participants to correctly select the term associated with each picture in the database. All 372 pictures (93 male front view, 93 male side view, 93 female front view, 93 female side view) were tested in one experimental block. The images were presented in random order, different for every observer. Stimuli were presented for 1 s. This presentation time was based on previous results, where identification accuracy for the same face stimuli was measured as a function of presentation time (Schmidtmann et al. (2016)). The presentation of the face image was followed by the presentation of the target (correct) term as well as three distractor terms. Importantly, in order to minimize a decision bias caused by specific terms, the distractor terms were randomly selected from the remaining 92 terms shown in Table 2. In other words, each observer was presented with different distractor terms for each face. The terms were presented on a mid-grey screen in a diamond-like arrangement (see Figure 1), corresponding to the cursor keys on a computer keyboard, which were used to by the observers to make their choice. The target term could occur in one out of four locations, which was randomly determined. The task for the observer was to choose the term most appropriate to the expression in the picture. Participants were given a break after each group of 93 presentations, i.e. three breaks in total.

**Figure 1.**
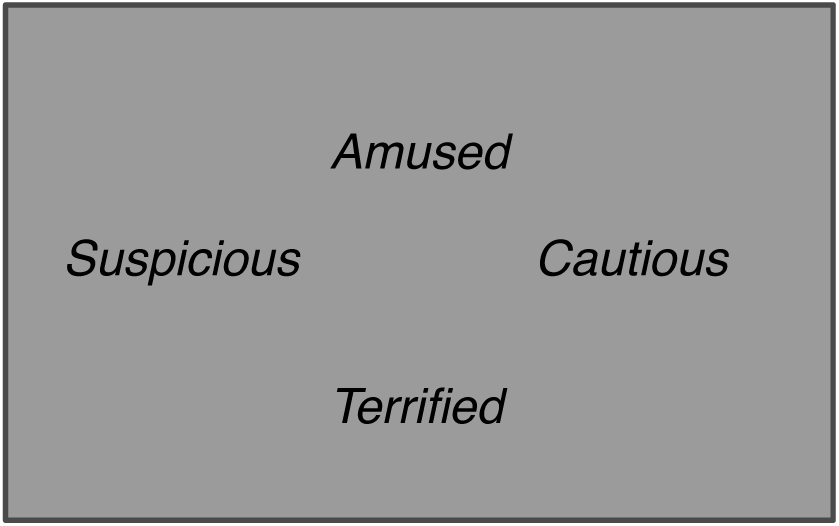
Experiment 1: Experimental Paradigm

### Results

Table 4 summarizes the performance (percent correct) across 33 subjects. The guess rate in a four-alternative forced choice paradigm is 25%. χ^2^-Tests with a Yates correction for continuity (*p* > .05) were performed to determine whether performances were significantly different from chance level for a given term (Yates (1934)). Performances not significantly better than chance are shown by the grey shading in Table 3 in the Appendix and by the lines in Figures 2 and 3 showing the sorted percent correct performances for the actors in front and side view as bar plots. Results show that for the pictures of the female actor, subjects performed significantly better than chance in 78 of 93 images (84%) for the front view condition and 74 of 93 images (80%) of the side view pictures. For the male actor, subjects performed significantly better than chance in 67 of 93 images (72%) in front view and 61 of 93 images (66% in side view. The non-significant terms are summarized in Table 4. Interestingly, 13 of these 52 non-significant cases occur in judgements of both the female and male actor. Furthermore, in 8 of these 52 terms subjects performed no better than chance for three or four of the images. These terms are indicated by the grey-shaded cells in Table 4.

**Table 3.**
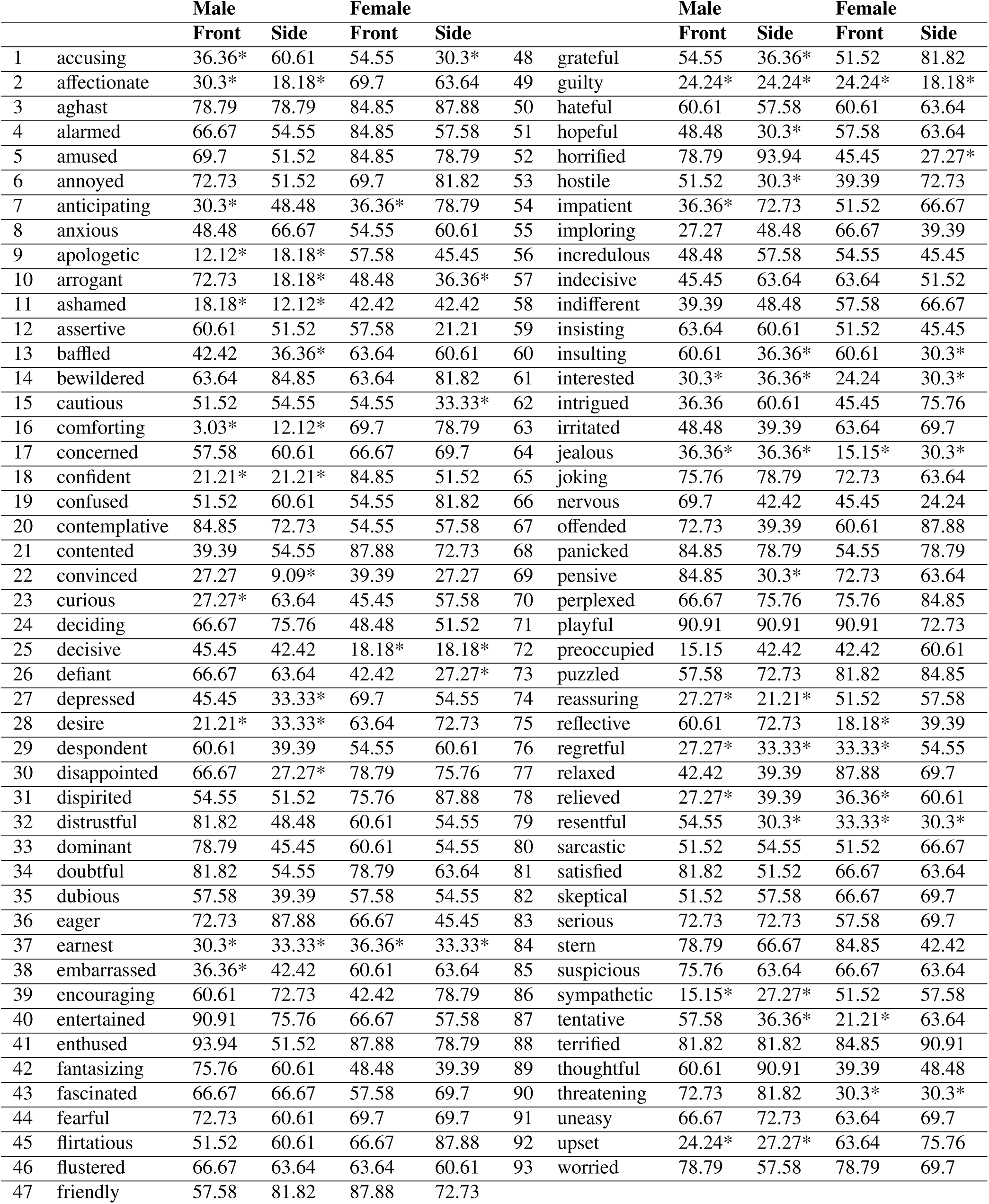
Percent correct for the images averaged across 32 subjects. The guess rate is 25%. Performances which are statistically not better than chance (χ^2^ – Yates correction for continuity; (α > .05) are indicated by the *.

**Table 4.** A summary of terms (sorted alphabetically) in which participants’ performances were not significantly better than chance. The cases which were not significant in three or more conditions are indicated by the *.

|  |  | **Female** |  | **Male** |  |  |  | **Female** |  | **Male** |  |
| --- | --- | --- | --- | --- | --- | --- | --- | --- | --- | --- | --- |
|  |  | **Front** | **Side** | **Front** | **Side** |  |  | **Front** | **Side** | **Front** | **Side** |
| 1 | accusing |  | 30.3 | 36.36 |  | 27 | hopeful |  |  |  | 30.3 |
| 2 | affectionate |  |  | 30.3 | 18.18 | 28 | horrified |  | 27.27 |  |  |
| 3 | anticipating | 36.36 |  | 30.3 |  | 29 | hostile | 39.39 |  |  | 30.3 |
| 4 | apologetic |  |  | 12.12 | 18.18 | 30 | impatient |  |  | 36.36 |  |
| 5 | arrogant |  | 36.36 |  | 18.18 | 31 | imploring |  | 39.39 | 27.27 |  |
| 6 | ashamed |  |  | 18.18 | 12.12 | 32 | indifferent |  |  | 39.39 |  |
| 7 | assertive |  | 21.21 |  |  | 33 | insulting |  | 30.3 |  | 36.36 |
| 8 | baffled |  |  |  | 36.36 | 34 | interested* | 24.24 | 30.3 | 30.3 | 36.36 |
| 9 | cautious |  | 33.33 |  |  | 35 | intrigued |  |  | 36.36 |  |
| 10 | comforting |  |  | 3.03 | 12.12 | 36 | irritated |  |  |  | 39.39 |
| 11 | confident |  |  | 21.21 | 21.21 | 37 | jealous* | 15.15 |  | 36.36 | 36.36 |
| 12 | contented |  |  | 39.39 |  | 38 | nervous |  | 24.24 |  |  |
| 13 | convinced* | 39.39 | 27.27 | 27.27 | 9.09 | 39 | offended |  |  |  | 39.39 |
| 14 | curious |  |  | 27.27 |  | 40 | pensive |  |  |  | 30.3 |
| 15 | decisive | 18.18 | 18.18 |  |  | 41 | preoccupied |  |  | 15.15 |  |
| 16 | defiant |  | 27.27 |  |  | 42 | reassuring |  |  | 27.27 | 21.21 |
| 17 | depressed |  |  |  | 33.33 | 43 | reflective | 18.18 | 39.39 |  |  |
| 18 | desire |  |  | 21.21 | 33.33 | 44 | regretful* | 33.33 |  | 27.27 | 33.33 |
| 19 | despondent |  |  |  | 39.39 | 45 | relaxed |  |  |  | 39.39 |
| 20 | disappointed |  |  |  | 27.27 | 46 | relieved* | 36.36 |  | 27.27 | 39.39 |
| 21 | dubious |  |  |  | 39.39 | 47 | resentful* | 33.33 | 30.3 |  | 30.3 |
| 22 | earnest* | 36.36 | 33.33 | 30.3 | 33.33 | 48 | sympathetic |  |  | 15.15 | 27.27 |
| 23 | embarrassed |  |  | 36.36 |  | 49 | tentative | 21.21 |  |  | 36.36 |
| 24 | fantasizing |  | 39.39 |  |  | 50 | thoughtful | 39.39 |  |  |  |
| 25 | grateful |  |  |  | 36.36 | 51 | threatening | 30.3 | 30.3 |  |  |
| 26 | guilty* | 24.24 | 18.18 | 24.24 | 24.24 | 52 | upset |  |  | 24.24 | 27.27 |

**Figure 2.**
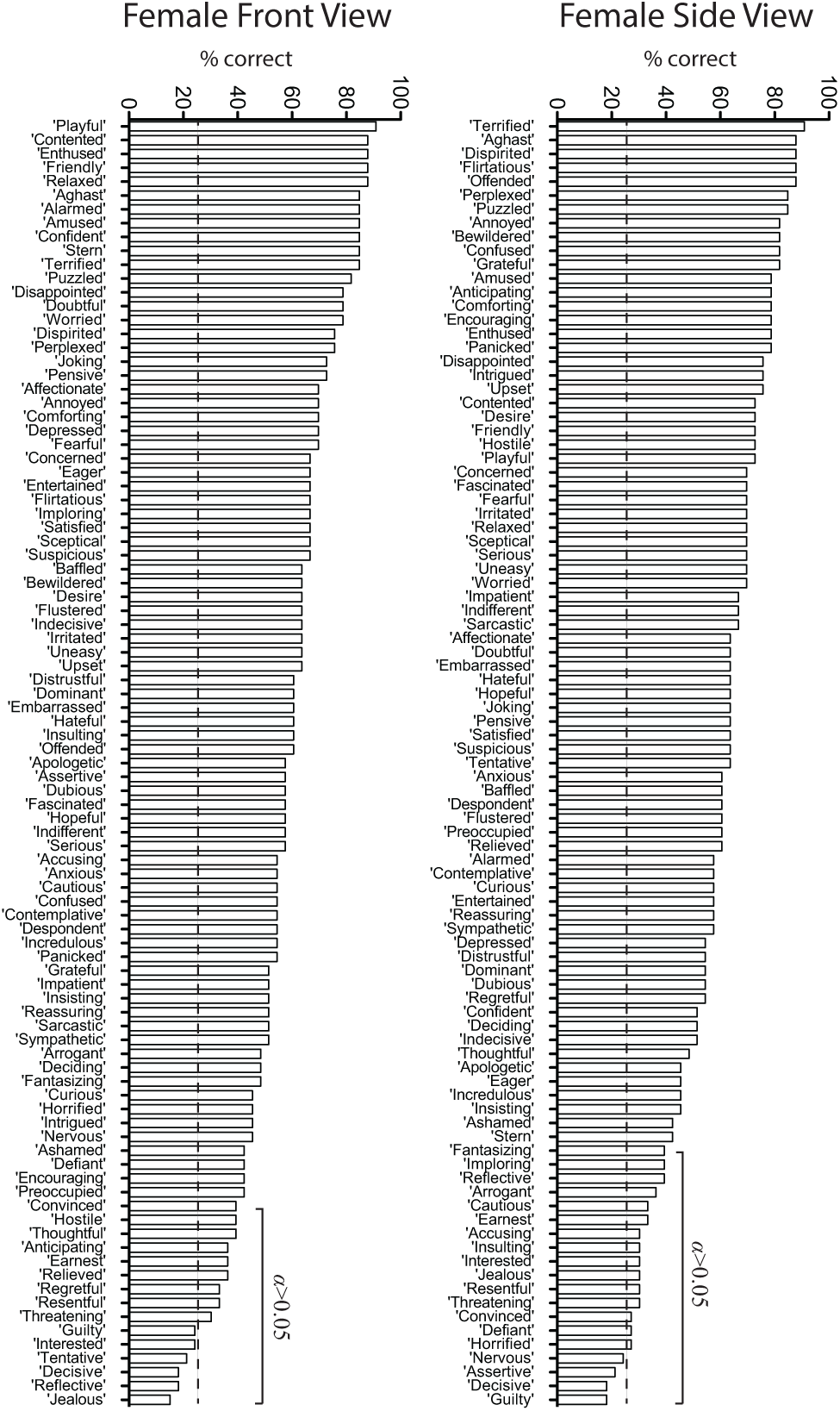
Bar plots showing percent correct for the 93 terms in the database for the female actor in both views. The dashed line represents the guessing rate (25 %). Performances which are statistically not better than chance (χ^2^ – Yates correction for continuity; α > .05) are indicated by the solid lines in each graph.

**Figure 3.**
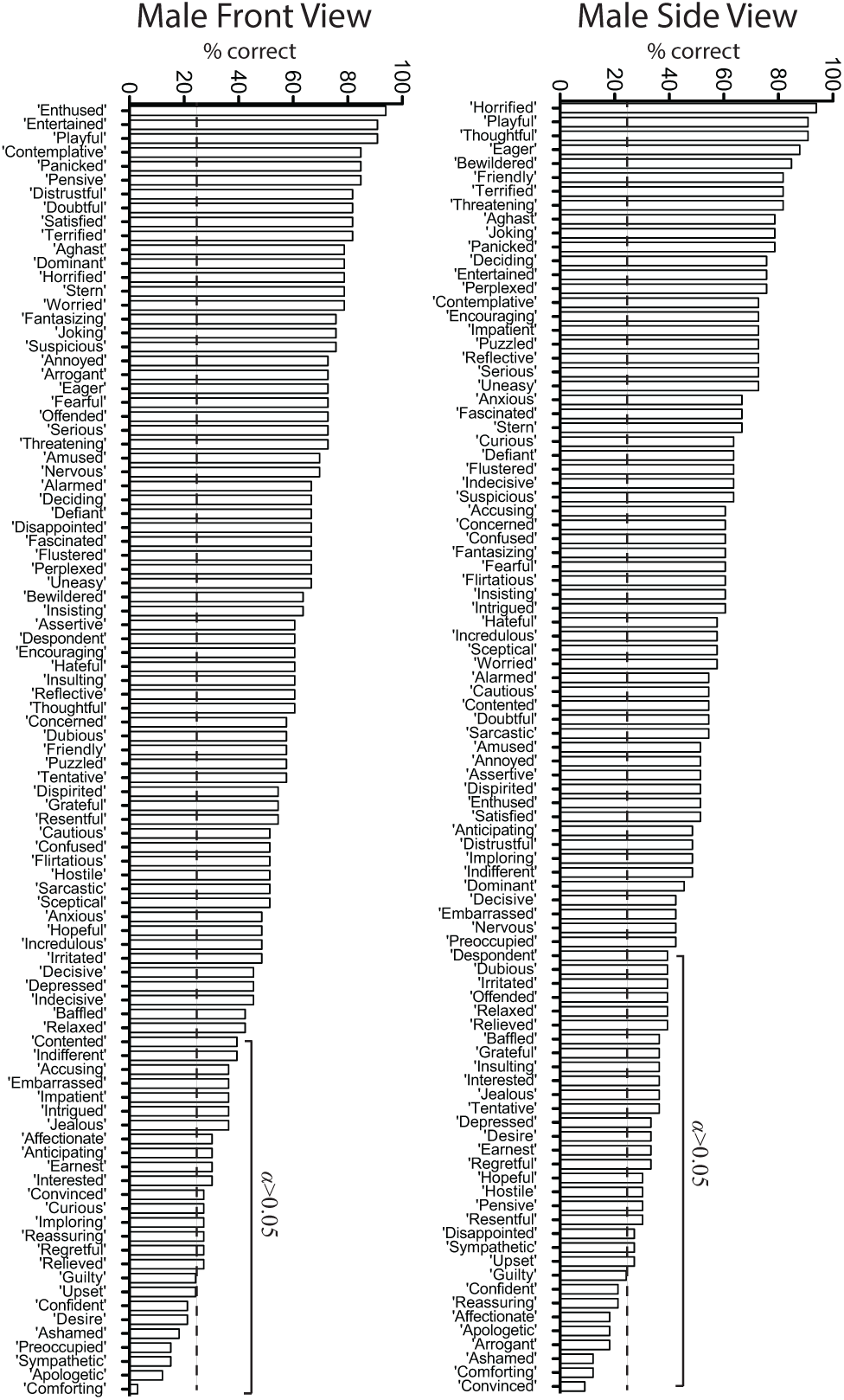
Bar plots showing percent correct for the 93 terms in the database for the male actor in both views. The dashed line represents the guessing rate (25 %). Performances which are statistically not better than chance (χ^2^ – Yates correction for continuity; α > .05 are indicated by the solid lines in each graph.

In addition, we conducted parametric Pearson correlation between each combination of the stimuli tested in experiment 1. Results show statistically significant correlations between results for the female faces in front and side view (*r* = .555, *p* < .001, *n* = 93), male faces in front and side view (*r* = .598, *p* < .001, *n* = 93), and female and male faces in front view (*r* = .336, *p* = .001, *n* = 93). All other correlations are presented in Table A1.

## Experiment 2

### Methods

#### Subjects

32 subjects participated in Experiment 2 (10 males, 22 females, mean age 22 years, ±4.13 SD).

#### Procedure

We employed a “point-and-click” task that did not rely on any semantic information being presented to observers during trials (Jennings et al. (2017)). The complete set of images (372) was presented in a random order. Each image was displayed for 1 s followed by the two-dimensional mental state-space (Russell (1980)), presented until the observer submitted a response (Figure 4 shows the 2-dimentional space). Once the two-dimensional space was displayed, the observers’ task was to click a computer mouse on the point within the space deemed most appropriate to the facial expression displayed in the image. The horizontal direction represented a rating of valence (pleasant vs. unpleasant) and the vertical direction a rating of arousal (low vs. high). Example emotions corresponding to different regions of the space are illustrated by the red text (not visible during testing) in Figure 4. The axes as well as the example mental states (red) were used to instruct the observer during training. In order to evaluate whether participants tended to locate facial expressions in similar regions of the two-dimensional space, we calculated an agreement score (η_*agreement*_) for each image among 32 observers in the following way.

**Figure 4.**
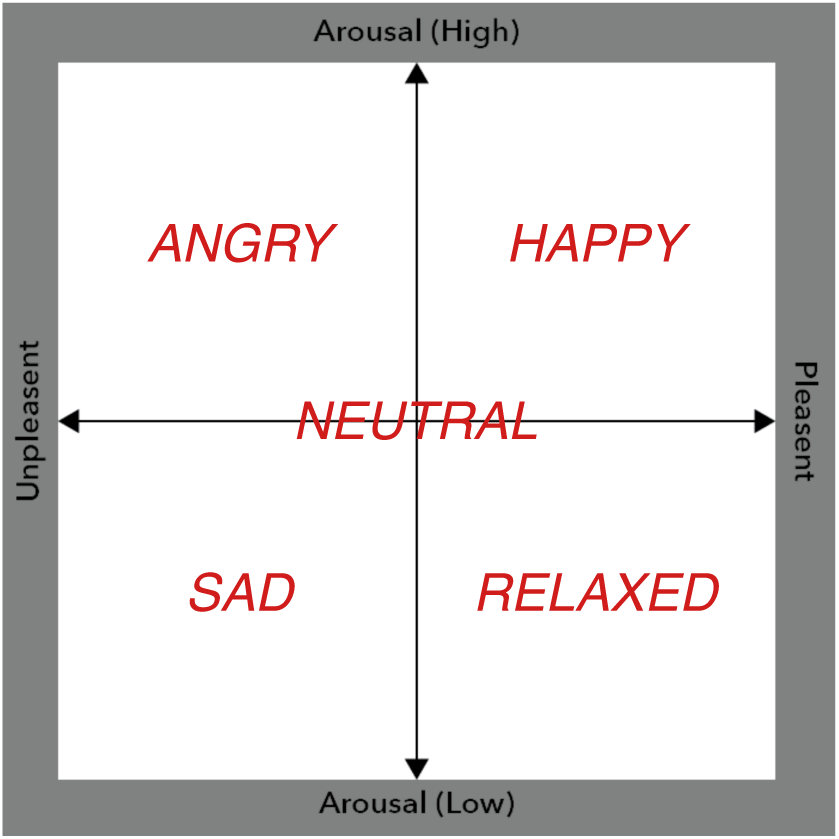
Experiment 2: The image was presented for 1 s, followed by the presentation of a valence-arousal space, extending from low to high arousal in one dimension and pleasant to unpleasant in the other dimension. Note: The red terms provide illustrations of the appropriate location of mental state terms used (the red text was not visible during testing

First, the mean arousal (*A*_*mean*_) and valence (*V*_*mean*_) coordinates were calculated across all observer responses for a given condition. Second, the Euclidian distance (r) for each of the observers’ response, and hence the mean rmean (see Eq. 1) was determined. Finally, these values were normalized (based on the highest mean value, *r*_*max*_) and shifted according to the lowest value (*r*_*min*_, see Eq. 2). This transformation produced agreement scores (η_*agreement*_) so that, a score of 1 corresponds to the greatest agreement between subjects and as the scores decrease the agreement between subjects’ decreases, i.e., emotion ratings were less tightly clustered around the mean location (see Eq. 3). Figure 5 illustrates the procedure for four hypothetical data points located within a subsection of the arousal-valence space.

**Figure 5.**
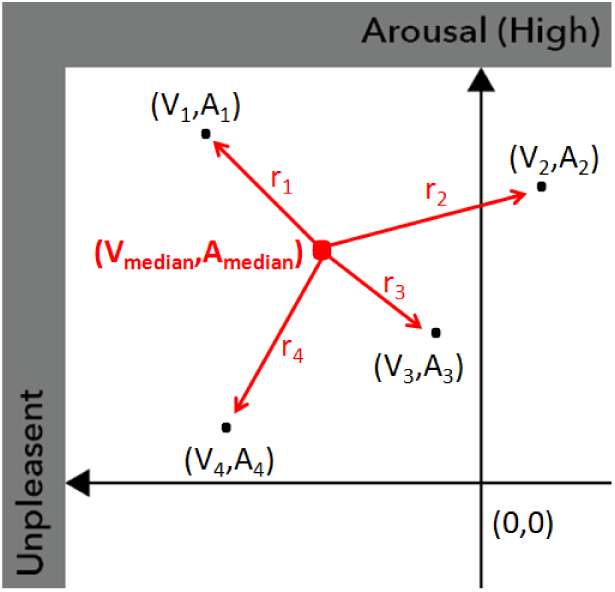
A subsection of the valence-arousal space showing four hypothetical responses (black dots); the red dot represents the mean valence (*V*_*mean*_) and arousal (*A*_*mean*_). The agreement score (η_*agreement*_) is determined by the mean Euclidian distance *r*.

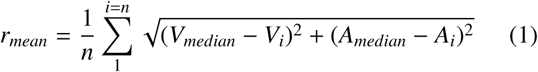

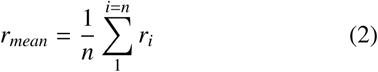

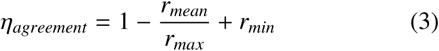

### Results

Tables 5 and 6 summarize the agreement scores (η_*agreement*_), for the female and male face stimuli, respectively. To visualize the magnitude of the agreement scores within the mental state-space three examples are illustrated in Figure 6. The circles are rendered with a radius equal to the values produced by Eq. 1 and the corresponding agreement values are stated for comparison. The results for each of the 93 terms can be downloaded here: McGill Face Database.

**Table 5.**
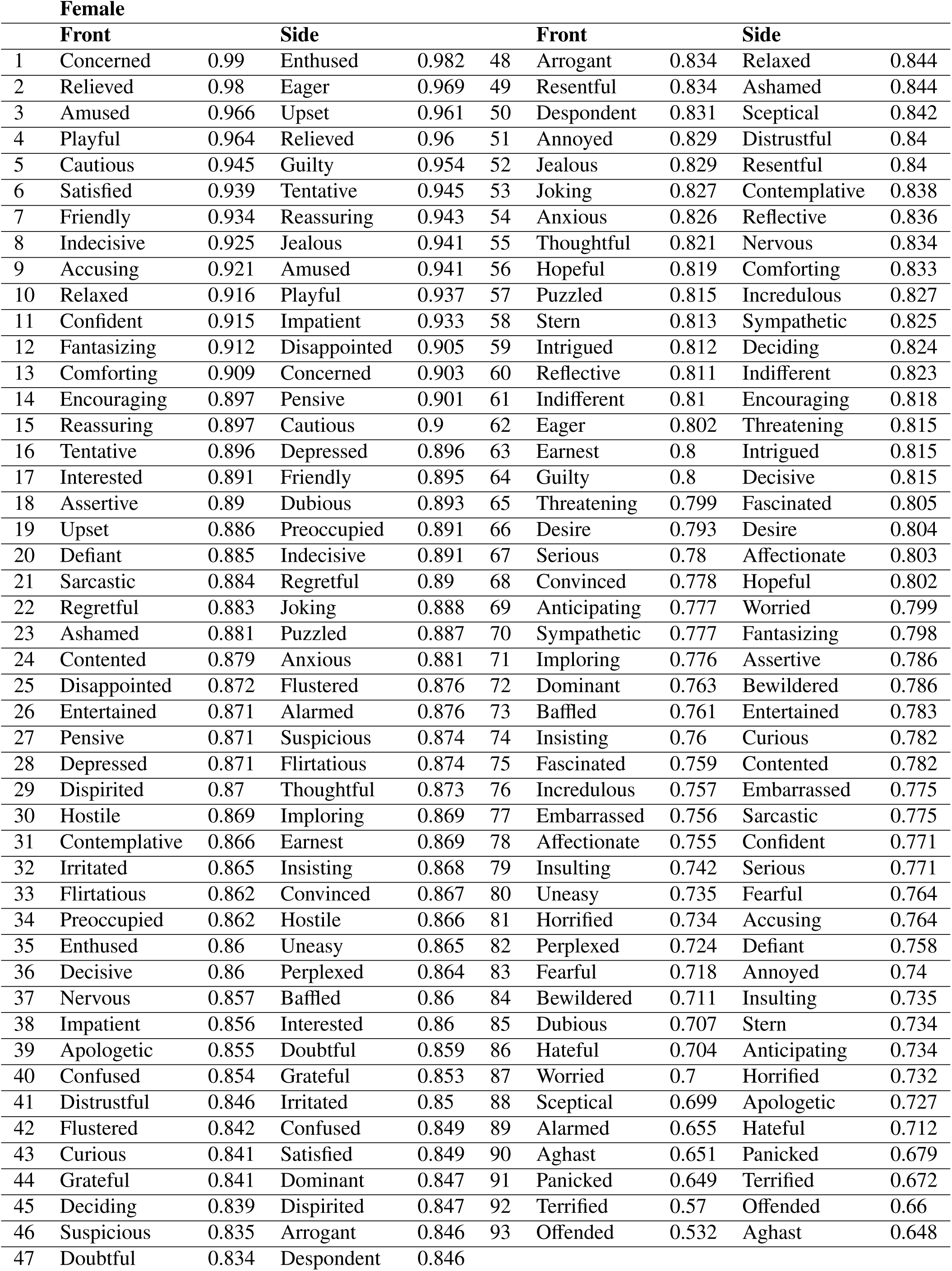
Agreement scores (η_agreement_) for the female in front and side view. Terms are sorted from high to low scores in each view.

**Table 6.**
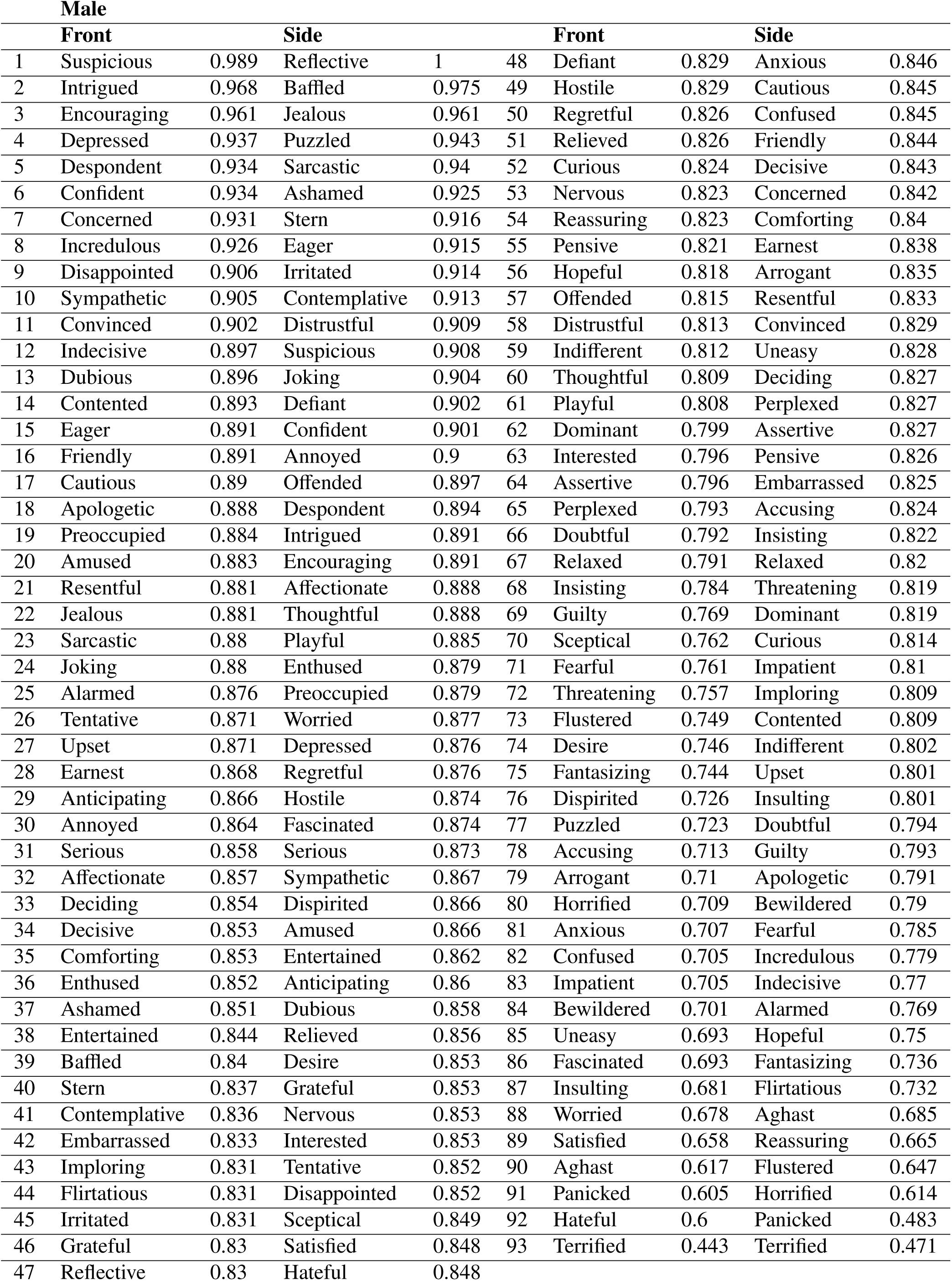
Agreement scores (η_agreement_) for the male actor in front and side view Terms are sorted from high to low scores in each view.

**Figure 6.**
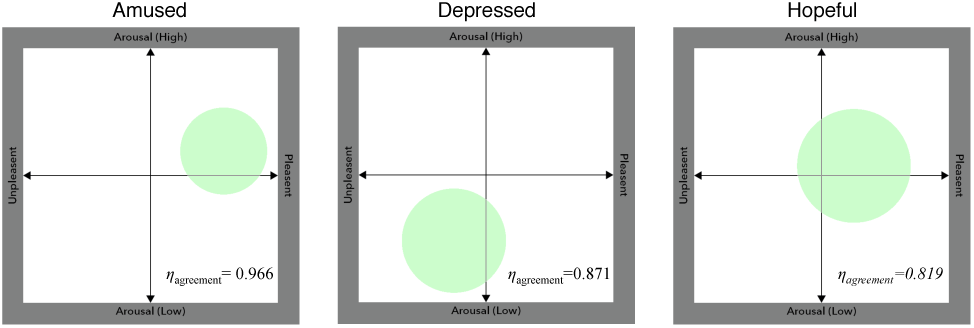
A subsection of the valence-arousal space showing four hypothetical responses (black dots); the red dot represents the mean valence (*V*_*mean*_) and arousal (*A*_*mean*_). The agreement score (η_*agreement*_) is determined by the mean Euclidian distance *r*.

### Correlations between Experiment 1 and Experiment 2

In a final analysis, parametric Pearson correlation tests were conducted between the percent correct performance for each stimulus in experiment 1 and the agreement score η_*agreement*_ for each stimulus in experiment 2. This analysis showed statically significant correlations between the results for male faces in front view in experiment 1 and male faces in front view in experiment 2 (*r* = -.302, *p* = .003, *n* = 93), for male faces in front view in experiment 1 and female faces in front view in experiment 2 (r = -.216, p = .038, n = 93) and for male faces in side view in experiment 1 and male faces in front view in experiment 2 (*r* = -.311, *p* = .002, *n* = 93) (see Table 7).

**Table 7.**
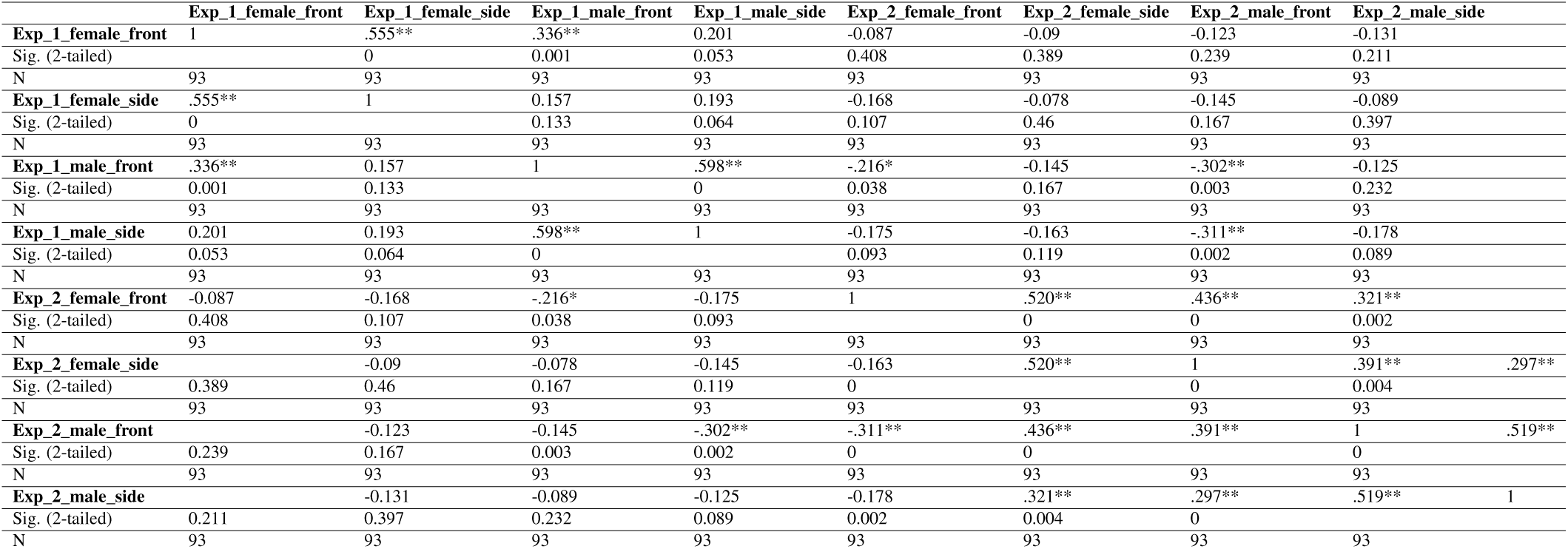
Parametric Pearson correlations / *. Correlation is significant at the 0.05 level (2-tailed). / **. Correlation is significant at the 0.01 level (2-tailed).

## Discussion

Most currently available image databases of facial expressions of mental states include only a very small range of possible mental states. With the exception of the “Mind Reading” platform (Baron-Cohen et al. (2004)), the vast majority of free databases employ the basic emotions proposed by Paul Ekman (e.g. Ekman, 1992): fear, disgust, surprise, happiness, sadness, and anger; see Table 1.) Even the full set of emotions, however, constitute only one category of mental state to which ToM is directed. In order to investigate ToM comprehensively, a more expansive set of stimuli is desirable. The aim of the current study was to develop and to validate a new database of such stimuli reflecting a greater variety of mental states. The McGill Face Database includes 4 representations of 93 mental state terms. The pictures are unmodified but can be altered if users wish to do so. In order to determine the usefulness of the database, two validation experiments were carried out. These experiments revealed considerable agreement among participants regarding the mental state expressed by the faces. Results from experiment 1 demonstrate that subjects can reliably select the correct term associated with a particular mental state despite the sematic complexity of the terms denoting them. Subjects performed significantly better than chance in 78 of 93 front view images and 74 of 93 side view images of the female actor, and they performed significantly better than chance in 67 of 93 front view and 61 of 93 side view images of the male actor. Results from this experiment also show that subjects performed better with images of the female actor, most likely because she was more expressive than the male actor. It is noteworthy that while subjects’ performance was better for front view images, the advantage over the side view was not dramatic (female: 84% vs. 80%; male: 72% vs. 66%). To our knowledge, this is the first demonstration of the high degree of accuracy human viewers exhibit when identifying complex mental states from only partially visible facial features. The Pearson correlation analyses for experiment 1 show a highly significant correlation between the two views of the same face as well as between front views of the male and female faces. The slightly more difficult side view task together with differences across the male and female faces presumably accounts for the absence of the full complement of correlations. The aim of the validation in experiment 2 was to develop a task that is independent of the complex vocabulary used in experiment 1. This approach has a number of advantages. First, some of the mental state terms may be more likely to be chosen just in virtue of their meanings. These biases would distort subjects’ performance. Secondly, the facial expressions produced by the actors are interpretations of mental state terms and some interpretations may be more easily associated with a target term than others. In this respect, the relationship between the facial expressions and the mental state terms explored in experiment 1 is distinctly different from the relationship between the basic emotions and the facial expressions to which they correspond. Whereas it is widely agreed that each basic emotion is represented by a single characteristic expression, many facial expressions might be thought to correspond to the mental state terms. Finally, it is of particular importance to be able to carry out ToM experiments without difficult vocabulary if one wants to study individuals with intellectual disabilities, or those suffering from conditions associated with impaired linguistic ability. The “point-and-click” paradigm in which subjects had to indicate the location of a given facial expression in a logical space (Russell (1980)), along the dimensions of valence and arousal, makes this possible (Jennings et al. (2017)). Results from this experiment show that there is substantial agreement across individuals about how to characterize faces along these dimensions. In addition, there is a high correlation between the face stimuli between perspectives and gender. The imperfect correlation between performance in the two experiments can be attributed to the presence of linguistic items in the first experiment and their absence in the second, as well as the difference in the specificity of the judgements required; the 2-dimensional space used in experiment 2 is a much coarser framework for classifying facial expressions than is the method of assigning a quite specific term to each face. The McGill Face Database thus provides a wide range of facial expressions of mental states that can be linked to mental state terms as well as accurately characterized in terms of arousal and valence independently of any such terms.

## Acknowledgement

We would like to thank Prof. Erin Hurley from the Department of English at McGill University for her help with the audition and recruitment process of the actors and for her helpful comments in that process. We further thank Scott Brevity Cope (photographer) and Edward Schokking for their help. This research was supported by a Social Sciences and Humanities Research Council of Canada grant #435-2017-1215, 2017 given to I.G. and G.S.

## Appendix

