## Supplementary Material for "The McGill Face Database: validation and insights into the recognition of facial expressions of complex mental states"

Raw data (n=32) is plotted for each of the 93 emotional states within the space employed in experiment 2, i.e., arousal: y-direction and valence: x-direction. The coloured points represent the following image conditions: Female (front view): black, Female (side view): red, Male (front view): green, Male (side view): blue.

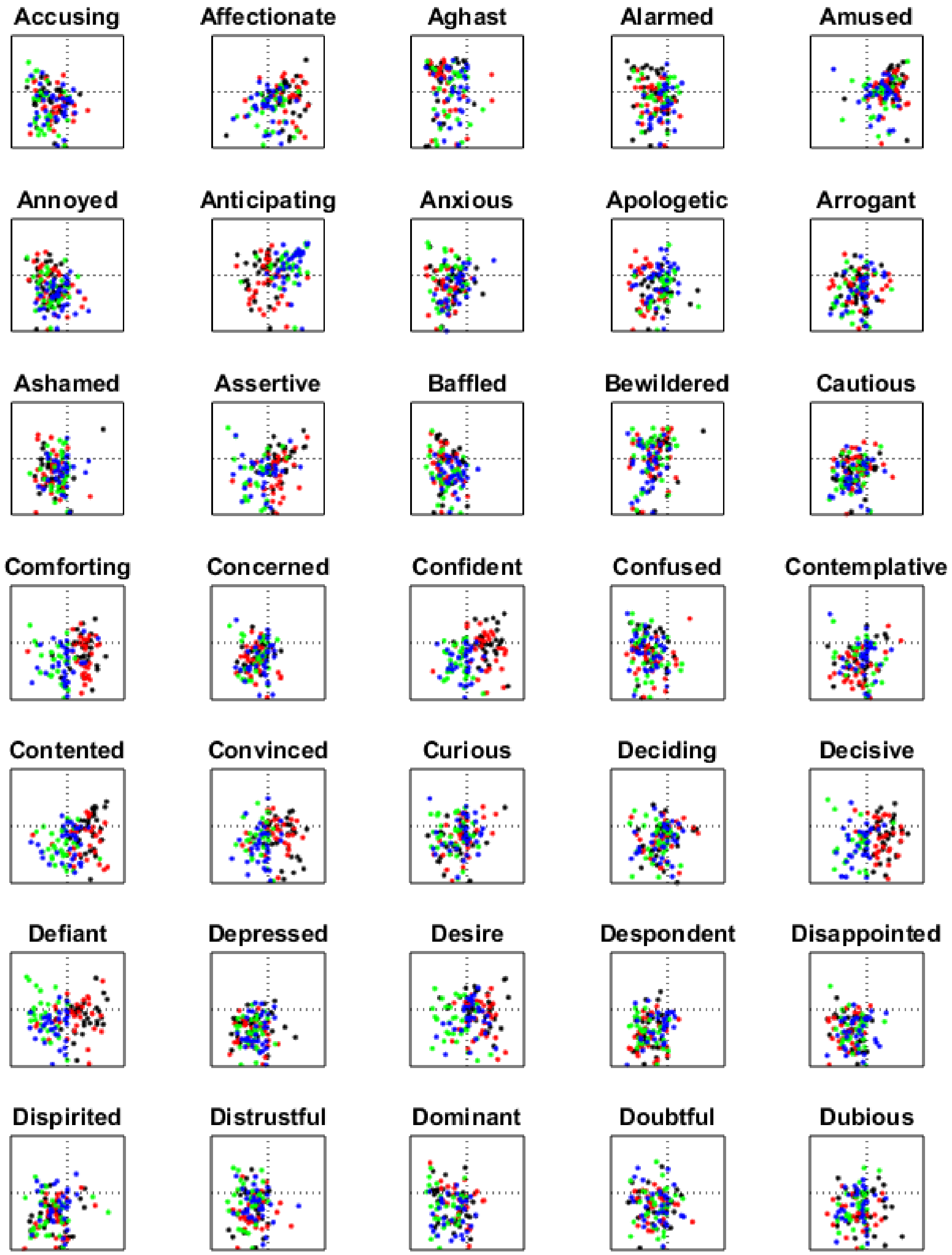

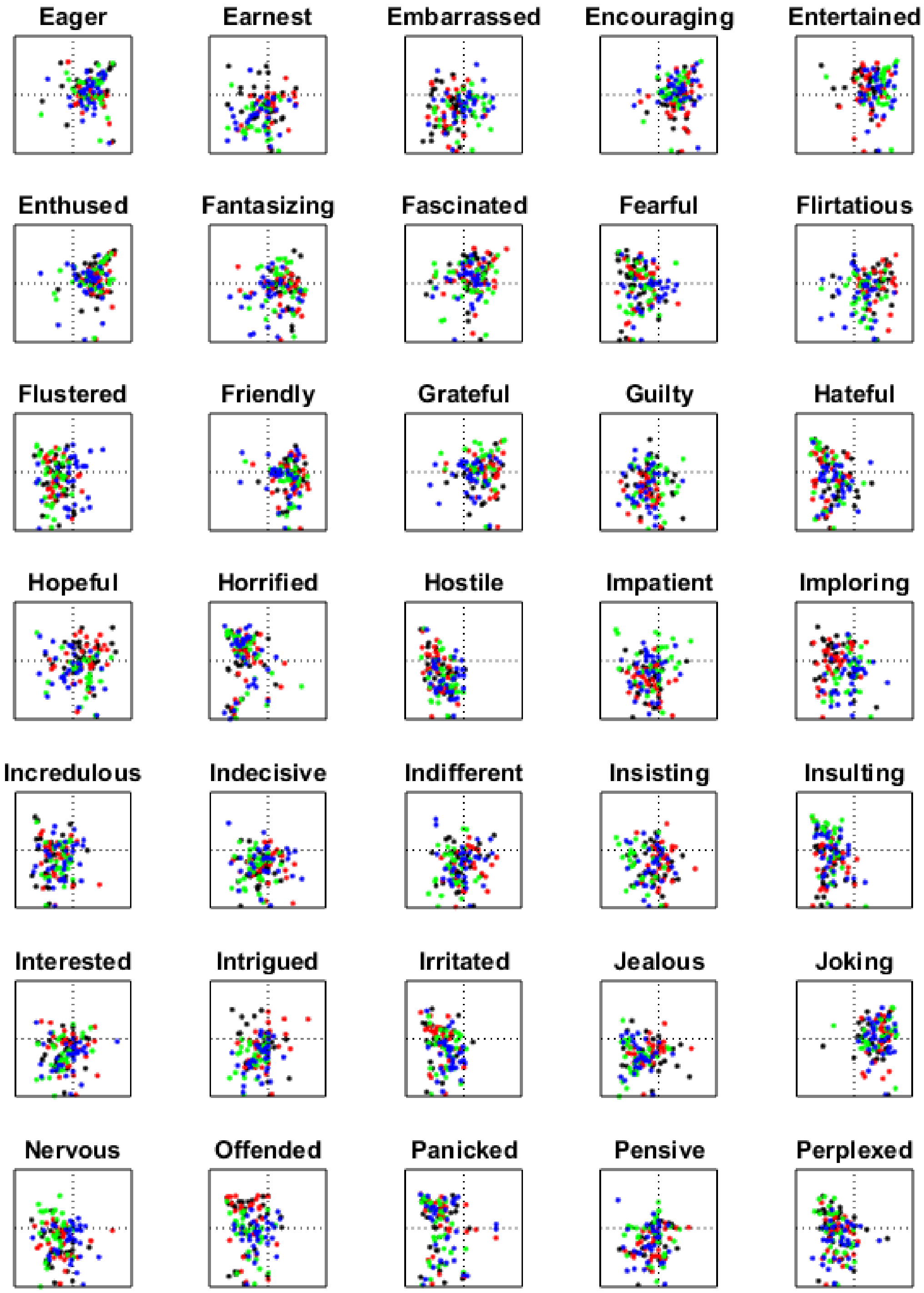
Eager
Earnest
Embarrassed
Encouraging
Entertained
Enthused
Fantasizing
Fascinated
Fearful
Flirtatious
Flustered
Friendly
Grateful
Guilty
Hateful
Hopeful
Horrified
Hostile
Impatient
Imploring
Incredulous
Indecisive
Indifferent
Insisting
Insulting
Interested
Intrigued
Irritated
Jealous
Joking
Nervous
Offended
Panicked
Pensive
Perplexed

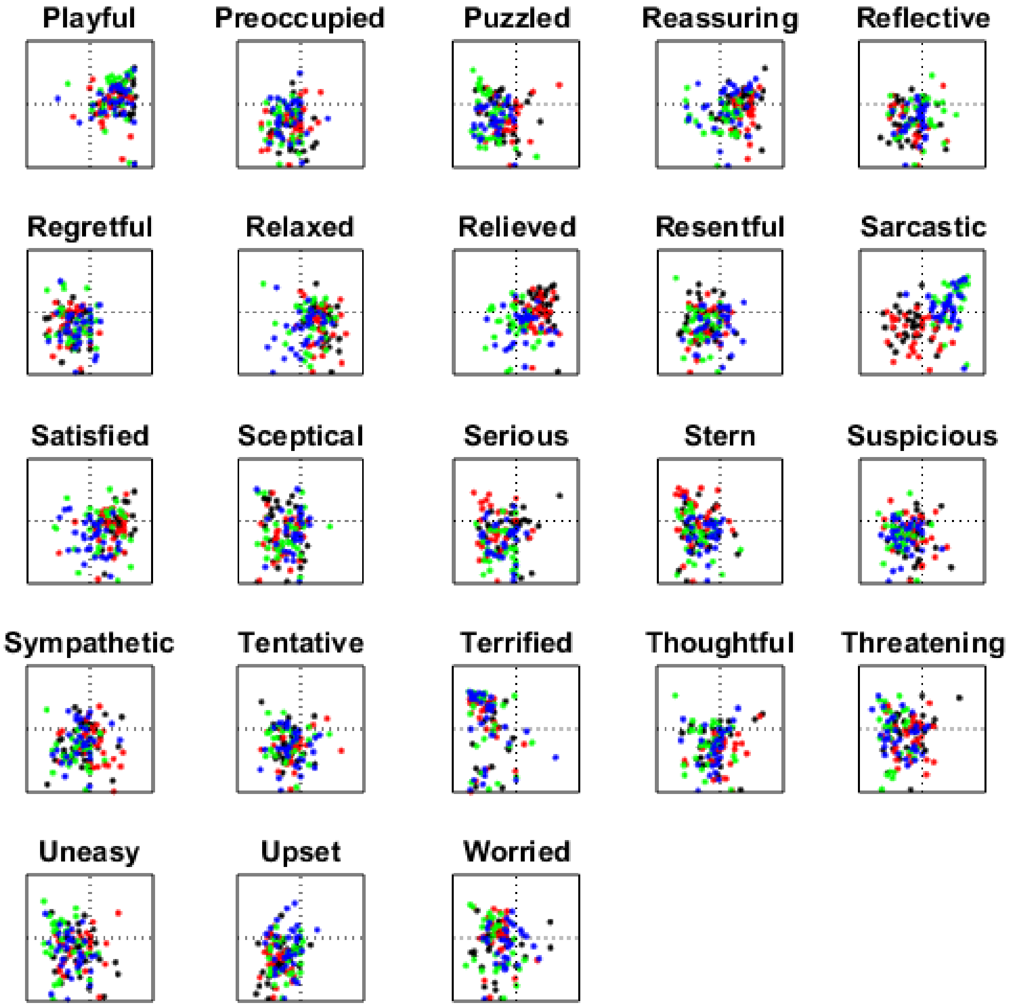
Playful
Preoccupied
Puzzled
Reassuring
Reflective
Regretful
Relaxed
Relieved
Resentful
Sarcastic
Satisfied
Sceptical
Serious
Stern
Suspicious
Sympathetic
Tentative
Terrified
Thoughtful
Threatening
Uneasy
Upset
Worried
